# Mutational escape from the polyclonal antibody response to SARS-CoV-2 infection is largely shaped by a single class of antibodies

**DOI:** 10.1101/2021.03.17.435863

**Authors:** Allison J. Greaney, Tyler N. Starr, Christopher O. Barnes, Yiska Weisblum, Fabian Schmidt, Marina Caskey, Christian Gaebler, Alice Cho, Marianna Agudelo, Shlomo Finkin, Zijun Wang, Daniel Poston, Frauke Muecksch, Theodora Hatziioannou, Paul D. Bieniasz, Davide F. Robbiani, Michel C. Nussenzweig, Pamela J. Bjorkman, Jesse D. Bloom

## Abstract

Monoclonal antibodies targeting a variety of epitopes have been isolated from individuals previously infected with SARS-CoV-2, but the relative contributions of these different antibody classes to the polyclonal response remains unclear. Here we use a yeast-display system to map all mutations to the viral spike receptor-binding domain (RBD) that escape binding by representatives of three potently neutralizing classes of anti-RBD antibodies with high-resolution structures. We compare the antibody-escape maps to similar maps for convalescent polyclonal plasma, including plasma from individuals from whom some of the antibodies were isolated. The plasma-escape maps most closely resemble those of a single class of antibodies that target an epitope on the RBD that includes site E484. Therefore, although the human immune system can produce antibodies that target diverse RBD epitopes, in practice the polyclonal response to infection is dominated by a single class of antibodies targeting an epitope that is already undergoing rapid evolution.

## Introduction

Control of the SARS-CoV-2 pandemic will depend on widespread population immunity acquired through infection or vaccination. But a little over a year into the pandemic, a proliferating number of new viral lineages are rising in frequency^1–6^. These emerging lineages have mutations at <1% of all residues in the viral spike, and at no more than 3 of the ∼200 residues in the spike receptor-binding domain (RBD)—yet these handfuls of mutations often substantially erode and in some cases even ablate the polyclonal neutralizing antibody response elicited by infection^7–16^.

A substantial fraction of the neutralizing activity of polyclonal antibody response to SARS-CoV-2 infection is due to antibodies that target the RBD^17–21^, although antibodies that target the NTD also contribute to neutralization^7–9, 22–24^. Structural and binding competition studies have shown that the most potently neutralizing anti-RBD antibodies target several distinct epitopes on the RBD’s receptor-binding motif^17, 19, 25–27^. However, the contributions of these different classes of RBD-targeting antibodies to the overall activity of the polyclonal antibody response remains less clear. It is therefore important to systematically determine both how viral mutations impact each antibody class, and how these antibody-specific effects shape the overall effects of viral mutations in a polyclonal context.

Here, we comprehensively map RBD mutations that reduce binding by structurally characterized representatives of three classes of neutralizing monoclonal antibodies that target the RBD’s receptor binding motif, as well as polyclonal plasma from convalescent individuals from whom some of the antibodies were isolated^21, 25, 28, 29^. We make these measurements by using a deep mutational scanning approach to systematically map how all RBD amino-acid mutations affect binding to yeast-displayed RBDs^30^ The resulting escape maps allow us to systematically compare how RBD mutations affect binding by the monoclonal antibodies, and we find that the antibodies cluster in the space of “viral escape” in a way that largely recapitulates prior classifications based on structural analyses of the antibody epitopes. However, some of the potently neutralizing monoclonal antibodies contribute very little to the escape maps of the polyclonal plasma, even for individuals from whom the antibodies were isolated. Instead, the plasma-escape maps usually most resemble a single antibody class (“class 2” in the Barnes et al. classification^25^) that targets the face of the receptor-binding ridge that is accessible in both “up” and “down” RBD conformations. Unfortunately, a mutation that escapes this antibody class (E484K) is present in many emerging viral lineages, including B.1.351, P.1, P.2, and B.1.526^1, 2, 4–6^. We suggest that domination of the RBD-targeting polyclonal response by a single antibody class is a major factor in enabling a small number of viral mutations to sometimes substantially erode neutralizing antibody immunity.

## Results

### Mapping all mutations that escape binding by key classes of RBD-targeting monoclonal antibodies

Most potent neutralizing antibodies against the SARS-CoV-2 RBD target the receptor-binding motif, where they compete for binding to ACE2^17, 25, 31^. Antibodies targeting the RBD have been divided into four major classes based on structural analyses of their epitopes; two with epitopes overlapping with the ACE2 binding site (class 1 and class 2), potent neutralizers that do not directly bind to the ACE2 contact surface (class 3), and antibodies that target a cryptic epitope outside of the receptor-binding motif and are generally less potent (class 4)^25^. We focused our studies on several antibodies representative of the first three classes of potently neutralizing receptor-binding motif-targeting antibodies. Class 1 antibodies bind the face of the receptor-binding motif that is accessible only when the RBD is in the “up” conformation (**Figure 1A**); the antibodies from this class in our study are C105 and LY-CoV016. Class 2 antibodies bind a face of the receptor-binding ridge that is accessible in both the “up” and “down” conformations (**Figure 1A**); the antibodies from this class in our study are C144, C002, and C121. Class 1 and 2 antibodies compete with ACE2 for RBD binding and have some overlap in their structural footprints particularly at ACE2 contact sites at the “top” of the receptor-binding ridge. Class 3 antibodies bind the opposite side of the receptor-binding motif (including the 443–450 loop), which like the class 2 epitope is accessible in both the “up” and “down” conformations, but has less overlap with the ACE2 binding footprint^25^ (**Figure 1A**); the antibodies from this class in our study are C135 and C110. All antibodies were isolated from humans previously infected with SARS-CoV-2^21, 28^ and have high-resolution structures^25, 28, 29^.

**Figure 1.**
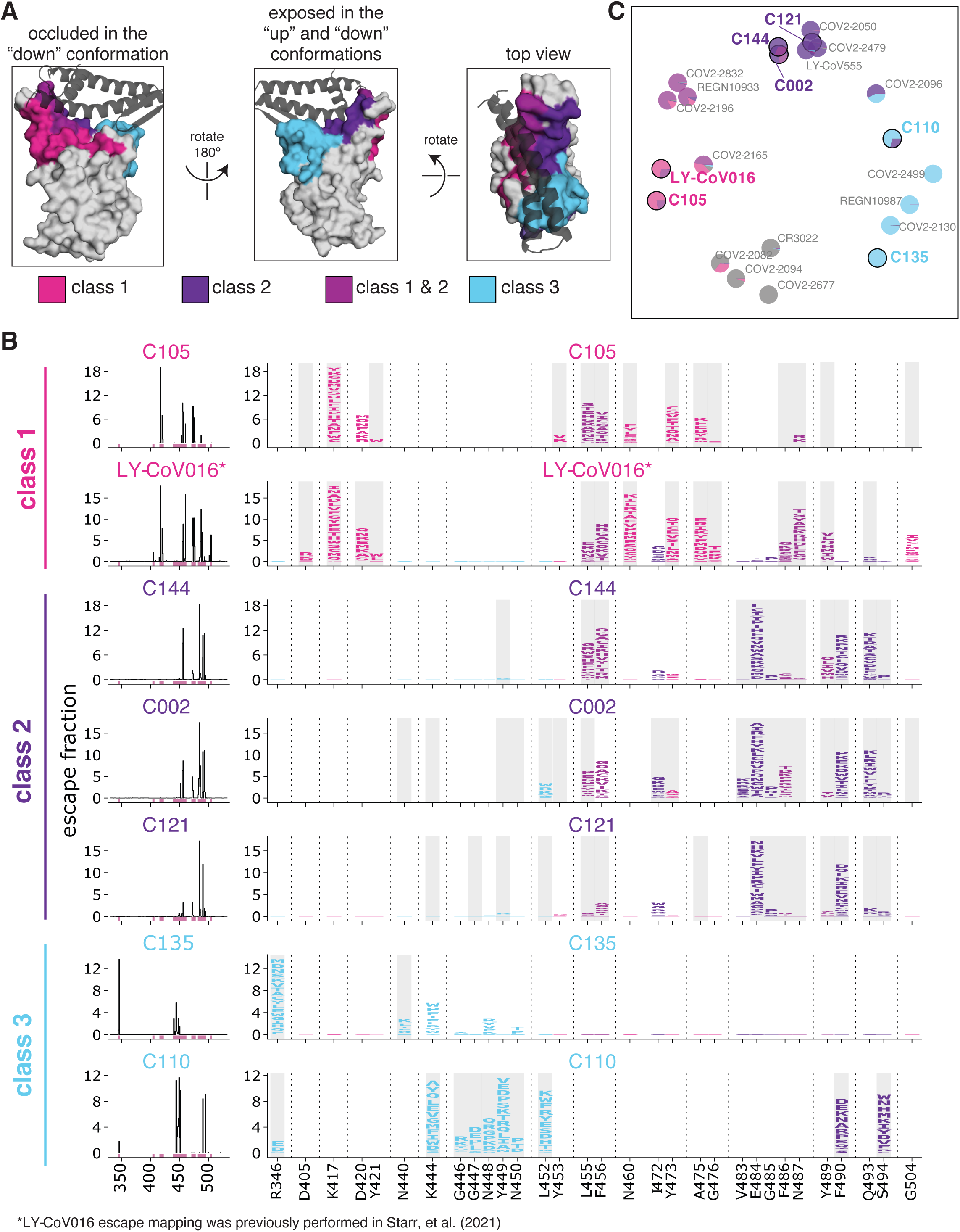
Maps of mutations to the RBD that escape binding by three classes of monoclonal antibodies that target the receptor-binding motif. **(A)** Epitopes for each of the three antibody classes^25^. ACE2 is shown as a gray cartoon. Some sites fall under both class 1 and class 2 and are shown as an intermediate pink-purple. **(B)** Escape maps for monoclonal antibodies from each of the three classes. The line plots at left indicate the summed effect of all mutations at each site in the RBD, with larger values indicating a greater reduction in antibody binding. The logo plots at right show the effects of individual mutations at key sites (indicated by purple highlighting on the x-axis of the line plots). In these logo plots, the height of each letter is that mutation’s escape fraction, so larger letters indicate mutations that cause a greater reduction in antibody binding. Sites in the logo plots are colored by RBD epitope. Sites that contact antibody (non-hydrogen atoms within 4Å in high-resolution structures) are highlighted with gray backgrounds. For C110, sites 444, 446, and 447 are unresolved in the structure but are likely in close contact with the antibody, and so are highlighted in gray. The data for LY-CoV016 were previously reported^33^ and are replotted here. **(C)** Multidimensional scaling projection of the escape maps, such that antibodies with similar escape mutations are drawn close together. Each antibody is represented as a pie chart colored according to the amount of escape in each RBD epitope. Antibodies for which escape maps are shown in panel **(B)** have black outlines and colored names. The other antibodies were profiled previously^30, 33, 35, 36^. Escape maps for all class 1, 2, and 3 antibodies in the plot are shown in **Figure S3**; for the class 4 antibody-escape maps, see Greaney et al. (2021)^30^. **Colors used in all panels**: sites are colored according to epitopes, as defined in Greaney et al. (2021)^25^. Bright pink for class 1, dark purple for class 2, medium pink-purple for class 1/2 overlap sites (455, 456, 486, 487, 489), cyan for class 3, and gray for all other RBD sites. Escape maps colored according to mutation effects on RBD expression and ACE2 binding are in **Figure S4**. The ACE2-bound RBD structure in (**A**) is from PDB 6M0J^58^.

We used a previously described yeast-display deep mutational scanning approach^30^ to map all amino-acid mutations to the RBD that escape binding by each antibody. Briefly, we incubated libraries of yeast expressing nearly all possible single amino-acid mutations in the RBD^32^ with each antibody, and then used fluorescence-activated cell sorting (FACS) to enrich for cells expressing RBD mutants that escaped antibody binding (**Figure S1**). We used deep sequencing to quantify each RBD mutation’s “escape fraction,” which is the fraction of cells with that mutation that fall into the FACS antibody-escape bin. These escape fractions range from 0 (no cells with the mutation fall into the antibody-escape bin) to 1 (all cells with the mutation fall into the antibody-escape bin). The escape fractions are well-correlated between independent libraries, and we report the average of duplicate measurements throughout (**Supplementary Table 1, Figure S2**). We represent the escape maps as logo plots, where the height of each letter is proportional to its escape fraction (**Figure 1B**). The escape map for LY-CoV016 has been previously described^33^; all other data using this assay were newly generated in this study. Interactive versions of these maps are available at https://jbloomlab.github.io/SARS-CoV-2-RBD_MAP_Rockefeller.

The escape maps show that antibodies within the same class are generally escaped by mutations at similar sites in the RBD, with large differences between classes (**Figure 1B**). For example, some class 1 but not class 2 antibodies were escaped by mutations to sites K417, D420, N460, and A475 (**Figure 1B**). Class 2 but not class 1 antibodies, on the other hand, were escaped by mutations at sites E484, F490, and Q493 (**Figure 1B**). Class 3 antibodies, which predominantly bind to the part of the receptor-binding motif opposite that of class 1 and class 2 antibodies (**Figure 1A**), tend to be escaped by a different set of mutations, including those at sites R346, K444, and G446–N450 (**Figure 1B**).

However, our maps emphasize that the antibody classes are approximate groupings, with overlap at some sites between classes. For example, class 1 and class 2 antibodies overlap in their binding footprints at the “top” of the receptor-binding ridge that contacts ACE2 (**Figure 1A**), and mutations to some of these overlapping sites (i.e., L455, F456, F486, and Y489) escape binding by some antibodies from both classes (**Figure 1B**). Similarly, the class 3 antibody C110 and the class 2 antibodies can both be escaped by mutations to site F490. There are also differences in escape mutations within antibody classes. For instance, the class 3 antibody C135 is strongly affected by mutations at site R346 but not L452, whereas the opposite is true for C110 (**Figure 1B**). Other examples can be found for the class 2 antibodies: C002 is escaped by some mutations to site L452, whereas C144 and C121 are less affected by these mutations (**Figure 1B**).

Our escape maps make it possible to group the antibodies using a more continuous approach, without strict divisions into three classes. Specifically, we used multidimensional scaling to project the escape maps into a two-dimensional space of binding escape (note that this computational technique has previously been used to visualize the antigenic relationships among strains of influenza virus^34^). We did this for the seven antibodies shown in **Figure 1B**, as well as 15 other antibodies for which we have previously determined escape maps^30, 33, 35, 36^, including class 4 antibodies, which bind to the RBD outside the receptor-binding motif and are generally less potently neutralizing^17, 19, 26, 31^. Antibodies with similar escape mutations are located close to one another in the multidimensional scaling projection, and antibodies with very distinct escape mutations are far apart (**Figure 1C**). The projection shows that while there is some clustering by structural class in space of escape, the antibodies are continuously distributed. For instance, the class 3 antibody C110 also selects escape at some “class 2 sites,” such as F490 and S494, and so C110 is located somewhat between the class 2 and 3 antibodies in the multidimensional scaling projection (**Figure 1C**).

### RBD mutations reduce antibody binding at only a subset of contact sites

We took advantage of the availability of high-resolution structures to compare the sites of escape mutations to the structural contacts between the antibodies and RBD. Most mutations that escape antibody binding are at sites in the RBD that directly contact the antibody; these sites are highlighted in gray in **Figure 1B**, with a structural contact defined as any non-hydrogen atom within 4Å. To visualize the escape mutations in a structural context, we mapped the extent of escape at each site to the structure of the antibody-bound RBD^25, 28, 29^ (**Figure 2A, S5A;** interactive, zoomable versions at https://jbloomlab.github.io/SARS-CoV-2-RBD_MAP_Rockefeller). All sites at which mutations strongly escape binding are in direct (<4Å) or proximal (4-8Å) contact with antibody in the resolved structures (**Figure 2B**).

**Figure 2.**
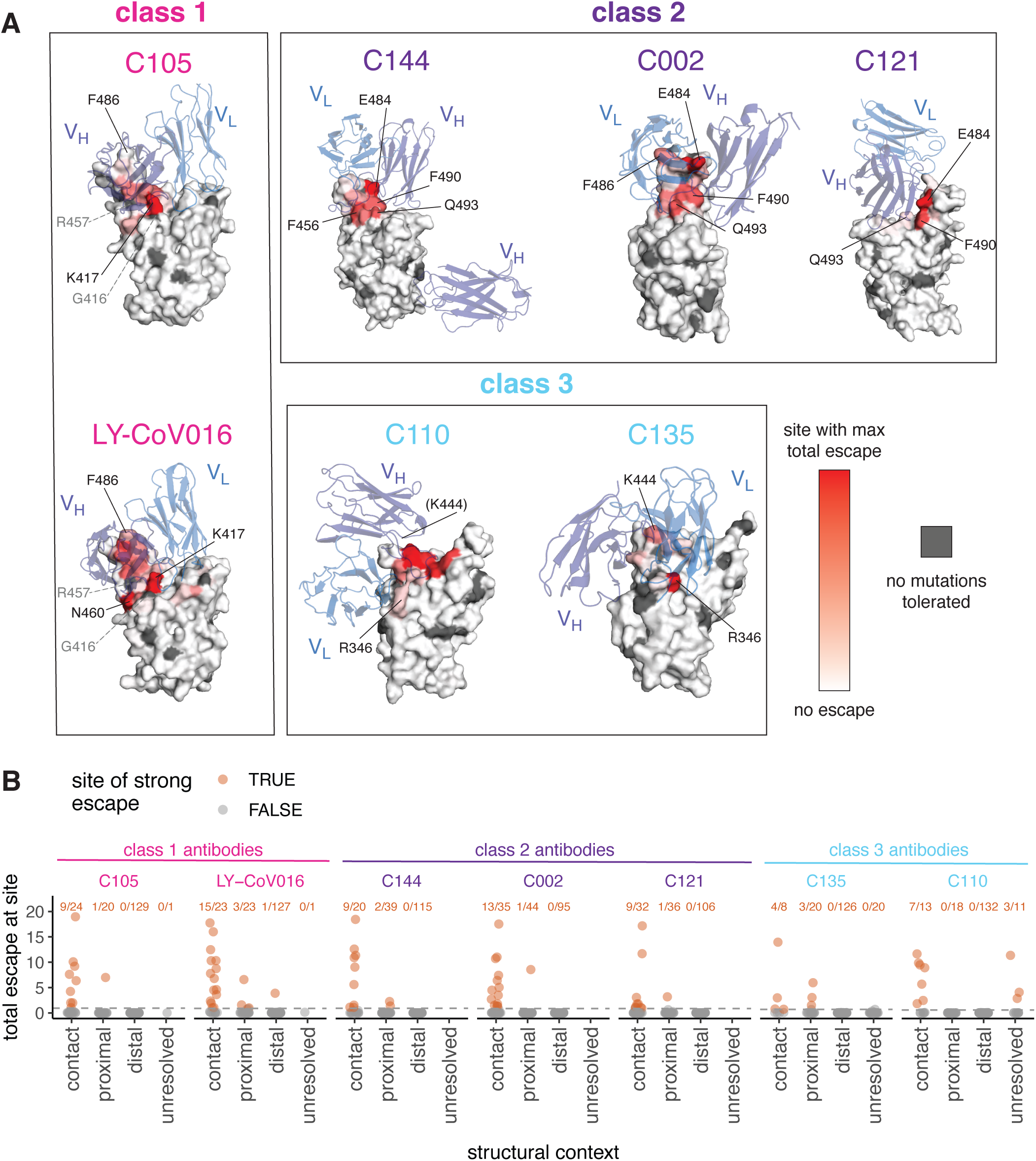
Mutations that escape antibody binding are usually in the direct structural footprint. **(A)** The total escape at each site is mapped onto the surface of the Fab-bound RBD, with white indicating no escape and red indicating the site with the most escape from that antibody. Sites where no mutations are tolerated for RBD folding or ACE2 binding are indicated in dark gray. For C105 and LY-CoV016, gray labels with dashed lines indicate example contact sites with no tolerated mutations. For C110, the general area where site 444 (unresolved in structure) would be located is indicated. **(B)** Total escape at each site in the RBD, with sites classified according to whether they are an antibody contact (within 4Å), antibody-proximal (4 to 8Å), antibody-distal (>8Å), or unresolved in the Fab-trimer structure. Text indicates the number of sites in each structural category that are sites of “strong escape,” (n / total) shown in orange. See **Methods** for details and PDB accessions.

However, not all antibody-contact sites had mutations that strongly escaped antibody binding (**Figure 2B, S5A**). There are several explanations. First, our approach maps functional antibody escape mutants that retain proper RBD folding and bind to ACE2 with ≥1% the affinity of the unmutated RBD^32^ (see **Methods**). Only 2,304 of the 3,819 possible amino-acid mutations to the RBD meet these criteria, and some sites have no tolerated mutations. For instance, G416 and R457 are both in the structural epitope of C105, but these sites have no tolerated mutations and thus do not appear in the escape map (sites with no tolerated mutations are indicated in dark gray in **Figure 2A, S5**). Second, sometimes mutations at antibody-contact sites simply do not strongly disrupt antibody binding^37^. For instance, site F486 is in structural contact with both C105 and LY-CoV016 and has many well-tolerated mutations, but mutations at this site more strongly affect the binding of LY-CoV016 than C105 (**Figure 1B, 2A, S5A**). Other examples include site R346, where nearly all mutations escape C135 but only charge-reversal mutations escape C110 (**Figure 1B, 2A, S5A**). Similarly, at site Q493, C144 and C002 are escaped by many mutations, but C121 is only escaped by Q493K/R (**Figure 1B, 2A, S5A**).

For some of the class 2 antibodies, the antibody makes a quaternary contact with an adjacent RBD in the context of spike trimer^25^ (**Figure S5B**). Our yeast-display system assays antibody binding to isolated RBD, and so does not map escape mutations to quaternary contact sites and cannot inform on their importance for antibody binding.

### The escape maps of polyclonal plasma often differ from those of monoclonal antibodies isolated from the same individual

The six antibodies newly mapped in this study were isolated from four different SARS-CoV-2 convalescent individuals (**Figure 3A**). Plasma was collected from these individuals at the same time that blood was collected for antibody isolation^21^. Because our escape-mutant mapping approach works for polyclonal sera or plasma in addition to monoclonal antibodies^18^, we mapped mutations that reduced binding by each of the four plasma plus one plasma sample without corresponding antibodies (**Figure 3B**; the plasma are prefixed with “COV-” to distinguish them from the antibodies which are prefixed with “C”).

**Figure 3.**
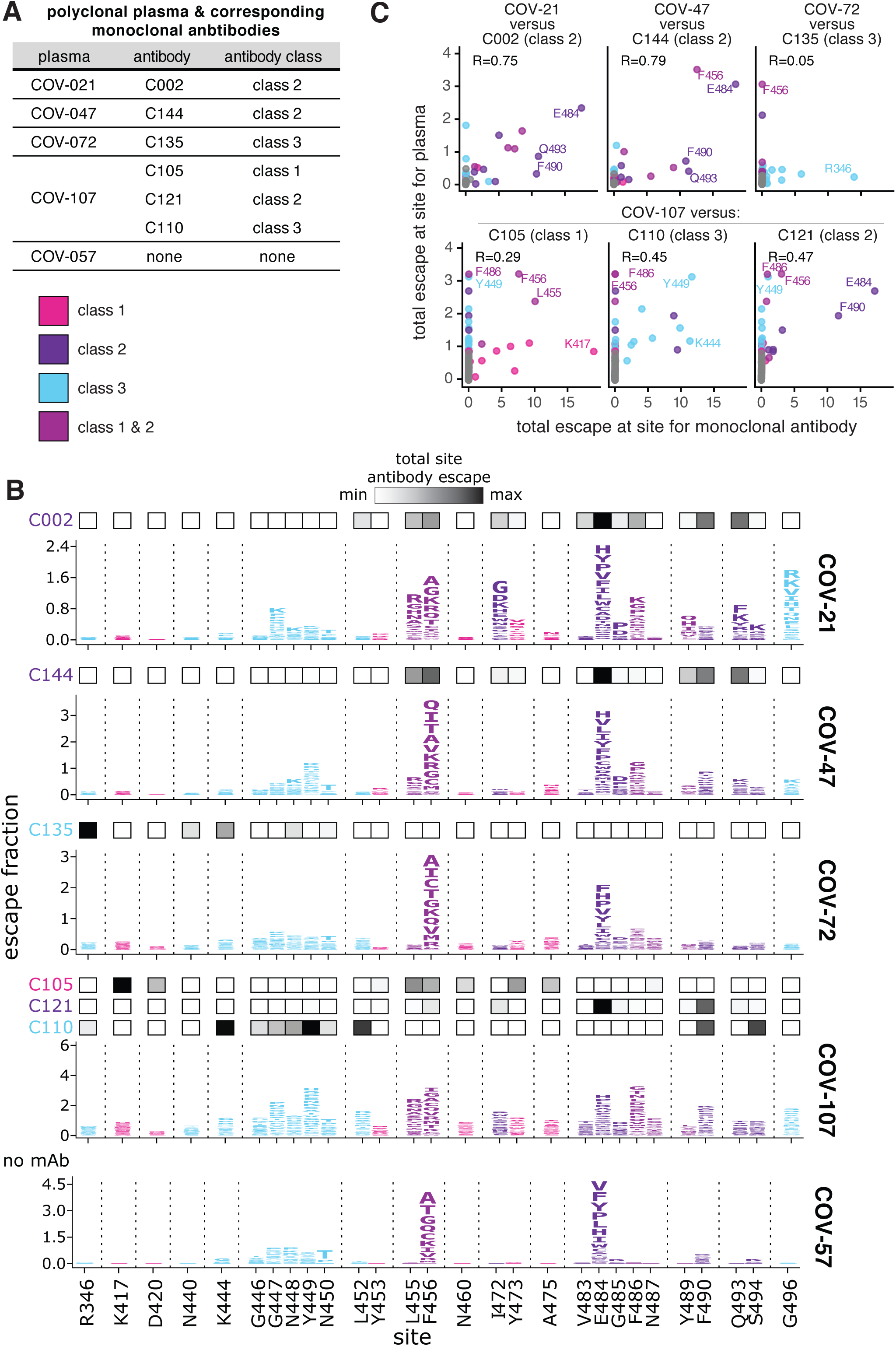
The mutations that reduce binding of polyclonal plasma often differ from those that reduce binding by monoclonal antibodies isolated from the same individual. **(A)** Table indicating which plasma and antibodies were derived from the same individual. **(B)** Escape maps for the polyclonal plasma antibodies, as in Figure 1B. The y-axis is scaled separately for each plasma (see **Methods**). When there are monoclonal antibodies isolated from the same individual, the total monoclonal antibody escape at each site is shown using the heat maps above the escape maps, with white indicating no effect and black indicating strong escape. **(C)** Correlation of plasma and monoclonal antibody escape for each plasma / antibody pair from the same individual. Each point in the scatter plots is a site, with the x-axis indicating the total escape at that site for the antibody and the y-axis indicating the total escape at that site for the plasma. Key sites are labeled. Pearson’s R shown above each plot. Colors in **B, C** reflect antibody classes as in Figure 1.

The plasma-escape maps shared many commonalities across all five individuals (**Figure 3B**), and generally resembled those from our prior study of a larger cohort of convalescent individuals^18^. In particular, mutations to sites F456 and E484 reduced binding for all five plasma samples (**Figure 3B**). Mutations to site E484 are of special note as the E484K mutation is present in the emerging B.1.351, P.1, P.2, and B.1.526 SARS-CoV-2 lineages^1, 2, 4–6^ and can reduce the neutralization titer of convalescent plasma by 3-fold or more^8, 10–12, 18^. Mutations to other sites, such as G446–N450 and F486 also reduce binding for some of the plasma samples profiled here (**Figure 3B**), consistent with our prior study of a larger cohort^18^. These findings suggest that while there is some heterogeneity among which mutations reduce binding of different individuals’ polyclonal plasma antibodies, there are also sites that are commonly targeted and should be monitored for antigenic evolution.

While the escape maps for the different plasma samples shared broad similarities, they often starkly differed from the escape maps of monoclonal antibodies isolated from the same individuals (**Figure 3B**, compare plasma escape maps in logo plots with overlay bars showing sites of escape for antibodies from the same individual). For instance, mutations to R346 had the largest effects on binding by the class 3 antibody C135, but had little effect on the same individual’s plasma (COV-72). Similarly, mutations at K417 had the largest effects on binding by the class 1 antibody C105, but had little effect on the corresponding plasma (COV-107). Conversely, mutations to site G496 reduced binding by the COV-21 plasma, but did not strongly affect any of the monoclonal antibodies in this study (**Figure 3B**). Overall, the correlations between the sites at which mutations escaped binding for the monoclonal antibodies and their corresponding polyclonal plasma were highest for the class 2 antibodies, and lower for the other antibody classes (**Figure 3C**).

### Class 2 antibodies contribute the most to the RBD escape maps of polyclonal plasma

To more broadly compare how antibodies of different classes contribute to the convalescent plasma escape maps, we used multidimensional scaling to project 22 antibodies and 28 polyclonal plasma into a two-dimensional space of binding-escape mutations (**Figure 4A**; the projection shows the 22 antibodies in **Figure 1C**, the 5 plasmas from **Figure 3**, and 23 plasmas from a previously characterized larger cohort^18^). The plasmas from both cohorts cluster together in the space of binding escape, but far from some of the antibodies (**Figure 4A**). In particular, most plasmas are positioned closest to the class 2 antibodies in the space of binding escape (**Figure 4A**).

**Figure 4.**
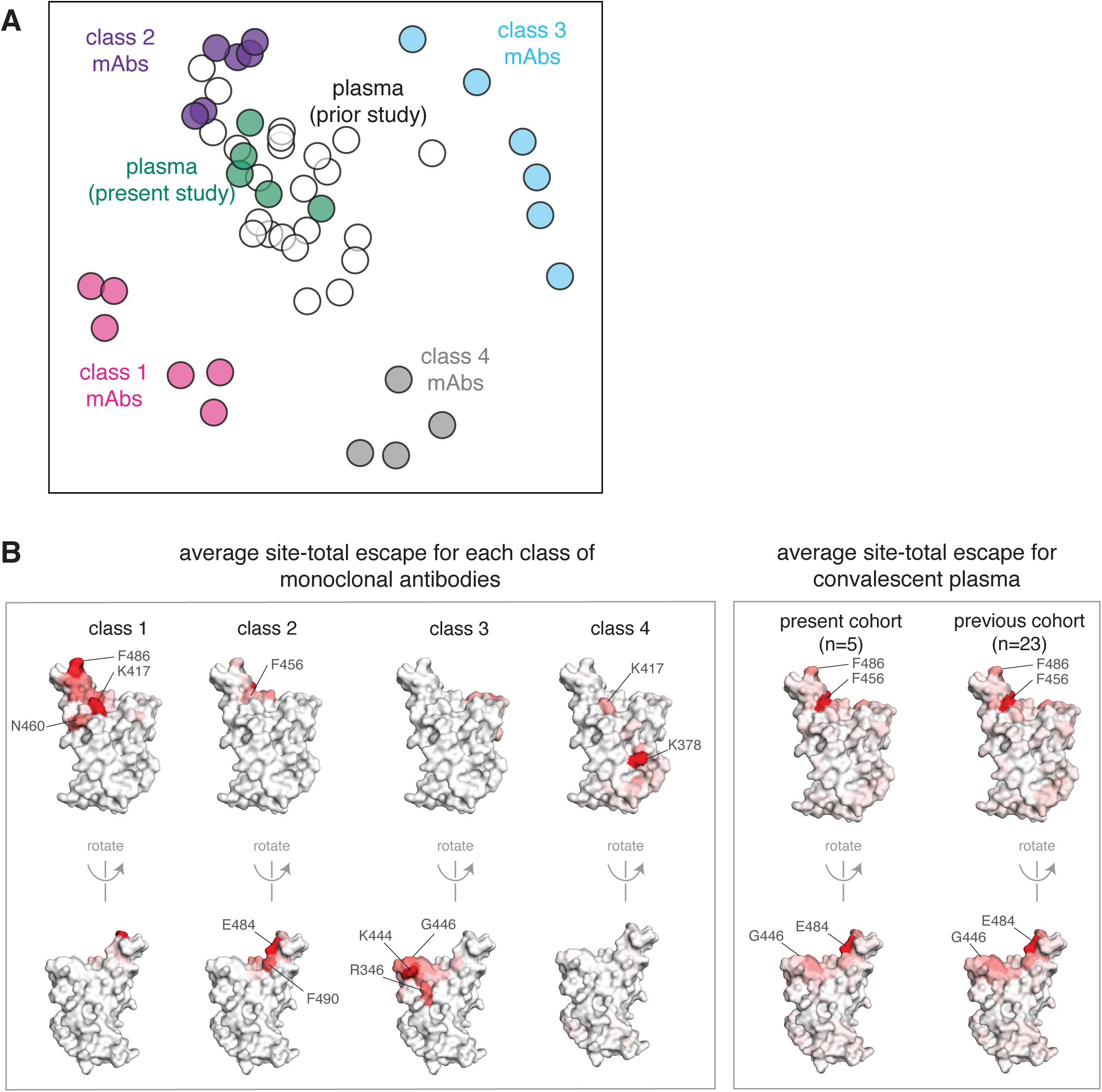
The escape maps of convalescent polyclonal plasma most resemble class 2 antibodies. **(A)** Multidimensional scaling projection of the escape maps of polyclonal plasma and monoclonal antibodies of each class. Antibodies or plasma that are nearby in the plot have their binding affected by similar RBD mutations. The antibodies are those in Figure 1C, colored according to antibody class. The five plasma newly mapped in this study are shown in green, and the previously mapped 23 plasma^18^ are shown in white. **(B)** Structural projection of sites where mutations reduce binding by each class of monoclonal antibodies (left) or polyclonal plasma (right). The RBD surface coloring is scaled from white to red, with white indicating no escape, and red indicating the site with the greatest average site-total escape for all antibodies or plasma in that group. Mutations to sites such as E484, F456, and F486 have some of the largest effects on binding by polyclonal plasma and class 2 antibodies.

To visualize the escape maps in terms of the RBD’s three-dimensional structure, we projected onto the surface of the RBD the total escape at each site averaged across all antibodies in a class, or all convalescent plasma in a cohort (**Figure 4B**). Again, the polyclonal plasma most closely resembled the class 2 antibodies. For instance, mutations to site E484 greatly reduced binding of both class 2 antibodies and polyclonal plasma (**Figure 4B**). Most of the class 1 antibody contributions to the plasma binding-escape maps came from sites shared with class 2 antibodies, such as F456 (**Figure 4B**), although note that mutations to F456 often do not strongly reduce plasma neutralization^18^. Consistent with the lesser contributions of class 1 antibodies, mutations at the class 1 site K417 had little effect on plasma binding, and others have found that the K417N mutation alone has a minimal-to-modest effect on plasma neutralization^10, 13, 15, 16^. Sites of escape from class 3 antibodies (e.g., G446) had visible effects on the polyclonal plasma, but again less so than for class 2 antibody sites (**Figure 4B**). However, in a prior larger study^18^, we found that mutations to the 443–450 loop strongly reduced binding of plasma from a minority of individuals, consistent with a few plasma falling closer to class 3 than class 2 antibodies in the space of binding-escape mutations (**Figure 4A**). Overall, these results show that class 2 antibodies usually dominate convalescent polyclonal plasma—although once a virus has accumulated mutations in class 2 epitopes (as has already occurred in some emerging lineages^1, 2, 4, 6^), then class 1 or 3 antibodies might dominate for the remaining anti-RBD antibody activity.

### Escape maps are consistent with the RBD mutations that arise when virus is grown in the presence of monoclonal antibodies

We assessed how well our escape maps predicted the actual antibody-escape mutations that arose when virus was grown in the presence of the antibodies. Prior work selected viral escape mutants by passaging chimeric VSV encoding the SARS-CoV-2 spike in the presence of several of the monoclonal antibodies we mapped in this study^7^. We hypothesized that the mutations selected during viral passage would reduce antibody binding without impairing ACE2 binding affinity. Accordingly, we examined how all of the selected mutations affected both antibody binding (as measured in the current study) and ACE2 affinity (as measured in our prior deep mutational scanning^32^). **Figure 5A,B** shows that in every case, the antibody-escape mutations selected in the virus were indeed among the single-nucleotide-change accessible amino-acid mutations that mediated the strongest escape from antibody binding without strongly impairing ACE2 affinity. Conversely, mutations that escaped antibody binding but were deleterious for ACE2 binding or RBD expression (e.g., E484V/A and mutations to sites 455 and 456) were not selected in the viral passaging. Therefore, our escape maps can be used in conjunction with prior data on the functional effects of RBD mutations to largely predict which escape mutations will arise when virus is grown in the presence of antibodies.

**Figure 5.**
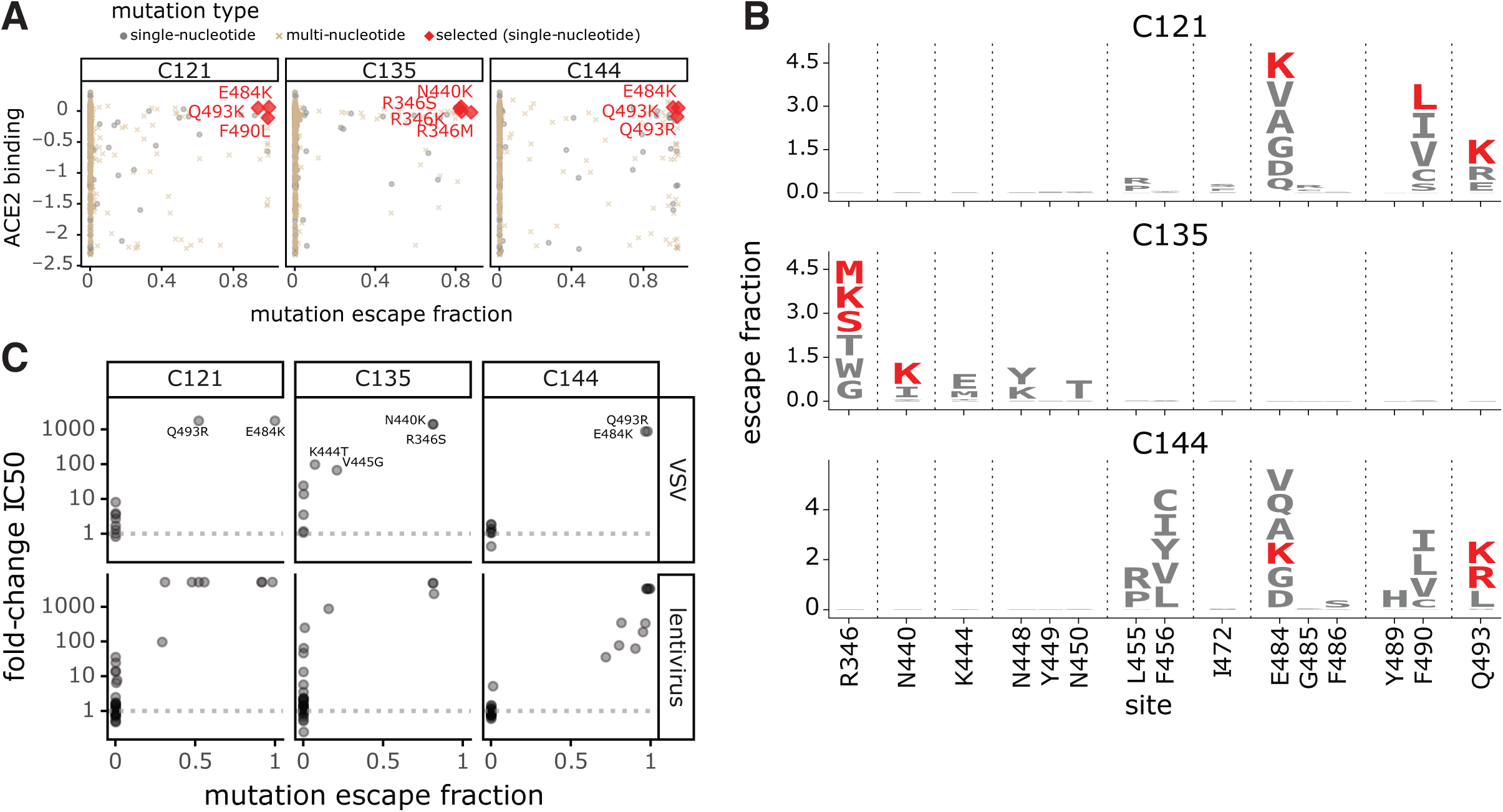
Escape maps predict mutations that are selected during viral growth in the presence of monoclonal antibodies. **(A)** Mutations selected when chimeric VSV encoding the SARS-CoV-2 spike was grown in the presence of each of the three indicated antibodies by Weisblum et al. (2020)^7^. Each point represents a different amino-acid mutation, with the x-axis indicating how strongly the mutation escapes antibody binding (measured in the current study) and the y-axis indicating how well the mutant binds to ACE2 (measured in Starr et al. (2020)^32^). The red diamonds indicate the mutations selected in VSV-spike by Weisblum et al., the gray circles indicate all other amino-acid mutations accessible by a single-nucleotide change, and the brown x’s indicate amino-acid mutations that require multiple nucleotide changes to the codon. **(B)** Logo plots showing the effects of only single-nucleotide accessible amino-acid mutations on antibody binding. Mutations selected in VSV-spike virus by Weisblum et al. are colored red. **(C)** The correlation of the effects of mutations on antibody binding measured in the current study and effects on viral neutralization previously measured by Weisblum et al.^7^ using chimeric VSV (top) or lentiviral particles (bottom). The x-axis shows the escape fraction measured in the current study, and the y-axis shows the fold-change in IC50 for viral neutralization caused by that mutation, such that larger numbers correspond to greater reductions in neutralization sensitivity. For effects of all antibody- and plasma-binding-escape mutations on ACE2 binding and RBD expression, see **Figure S4**. For each mutation’s escape fraction compared to fold-change IC50 against each monoclonal antibody or polyclonal plasma tested in Weisblum et al.^7^, see **Figure S6**.

Weisblum and colleagues also tested many RBD point mutations for their effects on neutralization of chimeric VSV or lentiviral particles by the antibodies studied here^7^. There was generally good agreement between our escape maps and these previously measured effects of mutations on viral neutralization. Nearly all mutations with a >100-fold reduction in neutralization also had large effects in our escape maps, although in a few cases mutations with more moderate effects on neutralization were not prominent in the escape maps (**Figure 5C, S6A**). Previously, we and others have reported that single point mutations can reduce the neutralization of some plasma by >10-fold, although other plasma are largely unaffected by any single mutation^7, 8, 18^. For the plasma in this study, prior work found that no tested mutation had such large effects on neutralization^7^. However, the class 2 antibody-escape mutation E484K did reduce neutralization by COV47 plasma by approximately 5-fold^7^, concordant with the prominence of site 484 in that plasma’s escape map (**Figure 3B, S6B**).

### Mutations that reduce binding by class 1 and 2 antibodies are present in emerging viral lineages

To assess the extent that SARS-CoV-2 has already acquired mutations that reduce binding by each antibody class, we compared the total escape at each site averaged across all antibodies of that class to the frequency of mutations at the site among sequenced SARS-CoV-2 isolates in GISAID as of March 15, 2021^38^. **Figure 6A** shows that mutations at sites targeted by each antibody class are present at appreciable frequencies among sequenced SARS-CoV-2 isolates. In particular, sites K417 and E484 are the strongest sites of escape for class 1 and class 2 antibodies, respectively—and mutations at both these sites are present in a substantial number of sequenced viral isolates (**Figure 6A**). In contrast, mutations that escape class 3 antibodies are currently not as prevalent among sequenced isolates (**Figure 6A**). While mutations have been observed at site K444 (the strongest site of escape from class 3 antibodies), these are at lower frequency than site 417 or 484 mutations (**Figure 6A**). Mutations at the class 3 site L452 are present at higher frequency, but L452 is only a moderate site of escape for this antibody class. As expected from the fact that class 2 antibodies dominate convalescent polyclonal plasma, the natural frequency versus escape plots for the plasma closely resemble those for class 2 antibodies, with site E484 clearly the most concerning mutation (**Figure 6B**).

**Figure 6.**
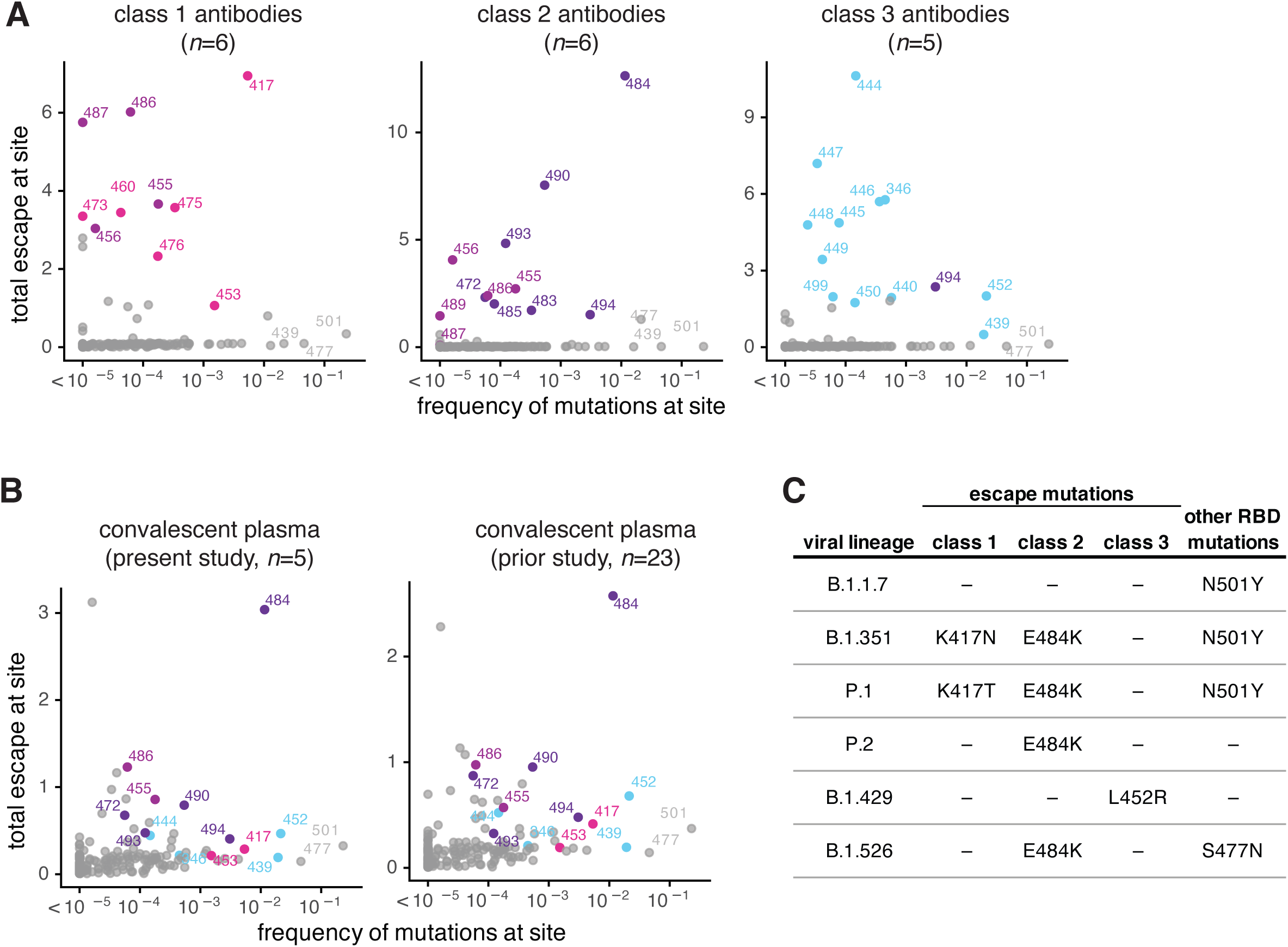
Mutations that escape binding by antibodies and plasma among sequenced SARS-CoV-2 isolates. **(A)** The total escape at each site averaged across antibodies in each class versus frequency of mutations at each site in GISAID sequences as of March 15, 2021. **(B)** The total escape at each site averaged across the polyclonal plasma versus frequency of mutations at each site in GISAID sequences. Left: plasma samples profiled in this study, right: plasma samples profiled previously^18^. **(C)** Antibody-escape mutations found in emerging viral lineages. RBD mutations in each lineage are assigned to the class of antibody they most strongly escape (e.g., E484K most strongly escapes class 2 antibodies but may also affect some class 1 antibodies). Other RBD mutations present in each viral lineage with negligible effects on binding of these antibodies are listed at right. For numbers of antibodies in each class, see *n* indicated in each panel in **A** and **B**. In each plot, key sites are labeled and colored according to RBD epitope using the same color scheme as in Figure 1.

We also examined the presence of escape mutations for each antibody class in some key emerging viral lineages (**Figure 6C**). All these emerging viral lineages except B.1.1.7^14, 16, 39–41^ have a mutation that escapes some antibodies from at least one class. The class 2 antibody escape mutation E484K is present in the B.1.351, P.1, P.2 and B.1.526 lineages (**Figure 6C**)^1, 2, 4–6^. Two of these lineages, and P.1, also have a class 1 antibody escape mutation, K417N or K417T, respectively (**Figure 6C**). Only the B.1.429 lineage carries a class 3 antibody escape mutation (L452R; **Figure 6C**). No viral lineages currently combine mutations that escape all three antibody classes—but the future emergence of a class 3 escape mutation in one of the lineages that already escape class 1 and 2 antibodies (B.1.351 or P.1) would be a worrying development, and should be monitored for closely.

## Discussion

We comprehensively mapped all mutations that escape binding by three major classes of antibodies targeting the SARS-CoV-2 RBD, and compared these escape maps to those for convalescent polyclonal plasma. We find that a single antibody class (class 2) largely shapes how RBD mutations affect binding by polyclonal plasma, even for individuals from whom potent neutralizing antibodies of other classes were isolated. The dominance of class 2 antibodies in RBD-targeting portion of polyclonal plasma could be due in part to the twin facts that their epitope is exposed in both “up” and “down” conformations of the RBD, and that such antibodies are often generated from frequently observed germline genes (including *VH1-2*, *VH3-53*, *VH1-69*)^21, 42–47,21, 48^. Consistent with our work here showing that class 2 antibodies shape how mutations affect polyclonal plasma binding, mutations at the site that most strongly affects binding by this antibody class (E484) have arisen multiple times in emerging viral lineages^1, 2, 4–6^. Therefore, our results show the importance of thinking about antigenic evolution in the context of different classes of antibodies that recognize different epitopes on the RBD.

Our work also sheds light on the extent to which it is functionally meaningful to subdivide anti-RBD antibodies into distinct classes based on their structurally defined epitopes. These structurally defined classes are inherently approximate groupings, since even antibodies with superficially similar structural epitopes bind to their antigens in subtly distinct ways^25^. Our comprehensive maps of binding escape mutations capture these subtle differences, and show that antibodies in the same structurally defined class can be differentially affected by the same mutation. Using the escape maps, we can visualize how the antibodies are related in terms of how their binding is functionally impacted by mutations at different RBD sites. These visualizations show that the arrangement of antibodies in the space of “viral escape” is indeed continuous, but that the class definitions based on structural analyses capture the high-level features of this arrangement since the structural footprint of an antibody largely determines which mutations most impact its binding. However, we suggest that in some cases, more continuous visualizations of the arrangements of antibodies in the space of viral escape such as the ones we present here could have benefits over structural classification schemes for analyzing the impacts of viral mutations.

Framing viral antigenic evolution in terms of different classes of antibody epitopes is also useful for interpreting the impacts of viral mutations and forecasting where in the RBD new escape mutations may arise in the future. Consistent with our work showing that mutations at class 2 antibody epitopes generally cause the largest reductions in the binding of polyclonal anti-RBD plasma antibodies elicited by infection, many emerging SARS-CoV-2 lineages have already acquired a mutation (E484K) at the site that most potently escapes antibodies of this class. Once viruses have escaped class 2 antibodies, antibodies of other classes (i.e., class 1 and 3) will contribute most remaining RBD-targeted antibody immunity. In this respect, it is noteworthy that two of the most prominent emerging viral lineages have also acquired a mutation (K417N/T) at the site that most potently escapes binding by class 1 antibodies. Fortunately, no viral lineages currently combine these class 1 and 2 escape mutations with a class 3 escape mutation, although a moderate class 3 escape mutation (L452R) is present in the B.1.427/B.1.429 viral lineage that lacks other RBD escape mutations^3, 49^. However, we suggest the appearance of a class 3 escape mutation on the background of a lineage that already has class 1 and 2 escape mutations such as K417N/T and E484K would be a worrying development, and should be monitored closely.

## Methods

### Data and Code Availability

- Complete computational pipeline for escape-mapping data analysis: https://github.com/jbloomlab/SARS-CoV-2-RBD_MAP_Rockefeller
- Markdown summaries of the escape-mapping data analysis steps: https://github.com/jbloomlab/SARS-CoV-2-RBD_MAP_Rockefeller/blob/main/results/summary/summary.md
- Raw data tables of mutant escape fractions are in **Supplementary Table 1**: https://github.com/jbloomlab/SARS-CoV-2-RBD_MAP_Rockefeller/blob/main/results/supp_data/all_samples_raw_data.csv.
- Raw Illumina sequencing for the escape mapping: NCBI SRA, BioProject: PRJNA639956, BioSample SAMN18148595.
- Processed Illumina sequencing counts for the escape mapping: https://github.com/jbloomlab/SARS-CoV-2-RBD_MAP_Rockefeller/blob/main/results/counts/variant_counts.csv

### SARS-CoV-2 convalescent human plasma samples

Plasma samples were previously described and collected as part of a prospective longitudinal cohort study of SARS-CoV-2 convalescent individuals in New York, NY^21^. Plasma samples profiled in this study were obtained 21-35 days post-symptom onset^21^. Samples were obtained upon written consent from community participants under protocols approved by the Institutional Review Board of the Rockefeller University (DRO-1006). All samples were heat-inactivated prior to use by treatment at 56 C for 60 minutes. Prior to use in each assay, plasma samples were centrifuged for 15 min at 2000 x*g* to pellet platelets.

### Neutralizing monoclonal antibodies binding the SARS-CoV-2 spike RBD

Antibodies were isolated from individuals in the cohort above as previously described^21^. Briefly, the IGH, IGL, and IGK genes were sequenced using IgG-specific primers from single-cell sorted RBD^+^, CD20^+^ memory B cells (CD3^−^CD8^−^CD14^−^CD16^−^CD20^+^Ova^−^RBD-PE^+^RBD-AF647^+^). Recombinant monoclonal antibodies were produced and purified as previously described^50, 51^. Antibodies were produced with a human IgG1 heavy chain and human IgK (C002, C110, C135) or human IgL2 (C105, C121, C144) constant regions. These antibodies were previously structurally characterized in complex with SARS-CoV-2 S trimer^25, 29^. A subset was functionally characterized in^7, 52^. The PBD accessions for the antibody-S complex structures are: 6XCM and 6XCN for C105, 7C01 for LY-CoV016, 7K8S and 7K8T for C002, 7K8X and 7K8Y for C121, 7K90 for C144, 7K8Z for C135, and 7K8V for C110^25, 28, 29^.

### RBD deep mutational scanning library

Monoclonal antibody and polyclonal clonal plasma selection experiments were performed in biological duplicate using a deep mutational scanning approach^30^ with previously described duplicate yeast-displayed mutant RBD libraries^32^. These libraries were generated in the RBD background of the SARS-CoV-2 isolate Wuhan-Hu-1 (Genbank accession number MN908947, residues N331-T531) via NNS codon tiling PCR mutagenesis, which introduced an average of 2.7 amino acid mutations per library variant. RBD variants were linked to unique 16-nucleotide barcode sequences to facilitate downstream sequencing and bottlenecked to library sizes of ∼100,000 uniquely barcoded variants. The libraries contain 3,804 of the 3,819 possible amino-acid mutations, with >95% present as single mutants on at least one barcode in the libraries. We previously used these libraries to measure the effect of all RBD mutations on yeast-surface RBD expression and ACE2 affinity^32^. As previously described, these libraries were sorted to eliminate variants that lose ACE2 binding prior to mapping the antibody-escape variants^30^.

### FACS sorting of yeast libraries to select mutants with reduced binding by polyclonal plasma

Antibody labeling and selection was performed essentially as described in Greaney, et al. (2020)^30^. Specifically, 9 OD aliquots of RBD libraries were thawed and grown overnight at 30°C 275 rpm in 45mL SD-CAA (6.7 g/L Yeast Nitrogen Base, 5.0 g/L Casamino acids, 1.065 g/L MES, and 2% w/v dextrose). Libraries were diluted to an OD of 0.67 in SG-CAA+0.1% dextrose (SD-CAA with 2% w/v galactose and 0.1% w/v dextrose in place of 2% dextrose), and incubated for 16-18 hours at room temperature with mild agitation to induce RBD surface expression. For each antibody selection, 20 OD units of induced cells were washed twice with PBS-BSA (0.2 mg/mL), and incubated in 4mL of PBS-BSA with monoclonal antibody or plasma for 1 h at room temperature with gentle agitation. Incubations were performed with 400 ng/mL for each monoclonal antibody (C105, C144, C002, C121, C135, or C110) or with a sub-saturating dilution of polyclonal plasma such that the amount of fluorescent signal due to plasma antibody binding to RBD was approximately equal across plasma (COV-021, 1:500; COV-047, 1:200; COV-057, 1:50; COV-072, 1:200; COV-107, 1:80). Labeled cells were washed with ice-cold PBS-BSA followed by secondary labeling for 1 h at 4°C in 2.5 mL 1:200 PE-conjugated goat anti-human-IgG (Jackson ImmunoResearch 109-115-098) to label for bound monoclonal antibody or 1:200 Alexa-647-conjugated goat anti-human-IgA+IgG+IgM (Jackson ImmunoResearch 109-605-064) to label for bound plasma antibodies, and 1:100 FITC-conjugated anti-Myc (Immunology Consultants Lab, CYMC-45F) to label for RBD surface expression. Labeled cells were washed twice with PBS-BSA and resuspended in 2.5mL PBS. Yeast expressing the unmutated SARS-CoV-2 RBD were prepared in parallel to library samples, labeled at the same 400 ng/mL and 100x reduced 4 ng/mL antibody concentrations for the monoclonal antibodies, and with 1x and 10x reduced plasma concentrations for the polyclonal plasma.

Yeast cells expressing RBD variants with substantially reduced antibody binding were selected via fluorescence-activated cell sorting (FACS) on a BD FACSAria II. For monoclonal antibody selections, FACS selection gates were drawn to capture 95% of yeast expressing unmutated SARS-CoV-2 RBD labeled at 100x reduced antibody concentration relative to library samples. For polyclonal plasma selections, FACS selection gates were drawn to capture 2.8–5% of the RBD mutants with the lowest amount of plasma binding for their degree of RBD expression (**Figure S1A-C**). Nearly zero (<0.1%) and 0.2 to 27.2% of cells expressing unmutated RBD fell into this gate when stained with 1x and 0.1x the concentration of plasma, respectively. For each sample, approximately 10 million RBD+ cells (range 8.7e6 to 1.5e7 cells) were processed on the cytometer, with between 1.5e6 and 2.0e6 monoclonal antibody-escaped cells and 3.2e5 and 5.3e5 plasma-escaped cells collected per sample. Antibody-escaped cells collected per sample into SD-CAA supplemented with 1% w/v BSA and grown overnight in 1.5mL SD-CAA + 100 U/mL penicillin + 100 µg/mL streptomycin at 30°C 275 rpm.

### DNA extraction and Illumina sequencing

Plasmid samples were prepared from up to 7.5 OD units (4e7 CFUs) of overnight cultures of antibody-escaped cells, and 30 OD units (1.6e8 CFUs) of pre-selection yeast populations (Zymoprep Yeast Plasmid Miniprep II) per manufacturer instructions, with the addition of a −80°C freeze-thaw step prior to cell lysis. The 16-nucleotide barcode sequences identifying each RBD variant were amplified by PCR and prepared for Illumina sequencing exactly as described previously^32^. Barcodes were sequenced via 50 bp single-end reads on an Illumina HiSeq 2500, targeting at least 2.5x as many sequencing reads as FACS-selected cells, and pre-sort reference populations of at least 2.5e7 reads.

### Analysis of mutant library deep sequencing and computation of per-mutant escape fractions

Escape fractions were computed as described in Greaney et al. (2021)^30^, with minor modifications as noted below. Specifically, we used the dms_variants package (https://jbloomlab.github.io/dms_variants/, version 0.8.5) to process Illumina sequences into counts of each barcoded RBD variant in each pre-sort and antibody-escape population using the barcode/RBD look-up table from Starr et al. (2020)^32^. Markdown renderings of these steps in the computational analysis are at https://github.com/jbloomlab/SARS-CoV-2-RBD_MAP_Rockefeller/blob/main/results/summary/aggregate_variant_counts.md and https://github.com/jbloomlab/SARS-CoV-2-RBD_MAP_Rockefeller/blob/main/results/summary/counts_to_cells_ratio.md.

For each antibody selection, we then computed the “escape fraction” for each barcoded variant using the deep sequencing counts for each variant in the original and antibody-escape populations and the total fraction of the library that escaped antibody binding via the formula provided in Greaney et al. (2021)^30^. These escape fractions represent the estimated fraction of cells expressing that specific variant that falls in the antibody escape bin, so a value of 0 means the variant is always bound by antibody and a value of 1 means that it always escapes antibody binding. We then applied a computational filter to remove variants with low sequencing counts or highly deleterious mutations that might cause antibody escape simply by leading to poor expression of properly folded RBD on the yeast cell surface. Specifically, we ignored all variants with pre-selection sequencing counts that were lower than the counts for the 99th percentile of the stop-codon containing variants because stop codon variants are largely purged by the earlier sorts for RBD expressing and ACE2-binding variants and so any residual presence provides an indication of low-count “noise.” Next, we removed any variants that had poor RBD expression or ACE2 binding, or contained mutations that individually cause poor RBD expression and ACE2 binding to eliminate misfolded or non-expressing RBDs. Specifically, we removed variants that had (or contained mutations with) ACE2 binding scores < −2.35 or expression scores < −1, using the variant- and mutation-level deep mutational scanning scores^32^. Note that these filtering criteria are slightly more stringent than those used in Greaney et al. (2021)^30^ but are identical to those used in Greaney et al. (2021) and Starr et al. (2021)^18, 33^. The ACE2 binding cutoff of −2.35 is used to represent the binding of RaTG13 to human ACE2^32^, which possesses the lowest known affinity capable of mediating cell entry^53^. The RBD expression cutoff of −1 is used to eliminate mutations that have as large an expression deficit as mutations to core disulfide residues. 2,034 of the 3,819 possible RBD amino acid mutations passed these filtering steps and were included in our escape maps. All previously reported escape mapping data^18, 30, 33, 36^ were reanalyzed in this study with the parameters listed above. A markdown rendering of the computation of the variant-level escape fractions and the variant filtering is at https://github.com/jbloomlab/SARS-CoV-2-RBD_MAP_Rockefeller/blob/main/results/summary/counts_to_scores.md.

Because some library variants contain multiple amino acid mutations, we next deconvolved variant-level escape scores into escape fraction estimates for single mutations using global epistasis models^54^ implemented in the dms_variants package, as detailed at (https://jbloomlab.github.io/dms_variants/dms_variants.globalepistasis.html). In this fitting, we excluded variants that contained mutations that were not seen as either single mutants or in at least two multiple-mutant variants. We then computed the estimated effect of each mutation as the impact of that mutation on the “observed phenotype” scale transformation of its “latent phenotype” as computed using the global epistasis models, and applied a floor of zero and a ceiling of 1 to these escape fractions. All of the above analysis steps were performed separately for each of the duplicate mutant libraries. We then only retained mutations that passed all of the above filtering and were measured in both libraries or had at least two-single mutant variant measurements in one library. The reported scores throughout the paper are the average across the libraries; these scores are also in **Supplementary Table 1**. Correlations in final single-mutant escape scores are shown in **Figure S2**. A markdown rendering of the computation that computes these mutation-level escape fractions is at https://github.com/jbloomlab/SARS-CoV-2-RBD_MAP_Rockefeller/blob/main/results/summary/scores_to_frac_escape.md.

For plotting and analyses that required identifying RBD sites of “strong escape” (e.g., choosing which sites to show in logo plots in **Figure 1A** or **Figure 3B** or label in **Figure 2B**), we considered a site to mediate strong escape if the total escape (sum of mutation-level escape fractions) for that site exceeded the median across sites by >10 fold, and was at least 10% of the maximum for any site. A markdown rendering of the identification of these sites of strong escape is at https://github.com/jbloomlab/SARS-CoV-2-RBD_MAP_Rockefeller/blob/main/results/summary/call_strong_escape_sites.md.

### Comparison of mutation escape fractions to previously measured neutralization concentrations

In **Figure 5C** and **Figure S6**, mutation-level antibody-escape fractions measured in this study are compared to previously measured neutralization titers (inhibitory concentration 50%, IC50) of the same monoclonal antibodies and polyclonal plasma against some RBD point-mutants^7^. The numerical IC50 values were extracted from figures in Weisblum et al. (2020)^7^ using the WebPlotDigitizer tool v4.4 (https://apps.automeris.io/wpd/). We convert those previously measured IC50s to fold-change IC50 relative to wildtype RBD. The numerical IC50s found in Weisblum et al. (2020)^7^ are also tabulated at https://github.com/jbloomlab/SARS-CoV-2-RBD_MAP_Rockefeller/blob/main/experimental_data/data/weisblum_ic50.csv.

### Analysis of mutations in circulating human SARS-CoV-2 strains

For the analysis in **Figure 6**, all 765,455 spike sequences on GISAID^38^ as of March 15, 2021 were downloaded and aligned via mafft^55^. Sequences from non-human origins and sequences containing gap or ambiguous characters were removed, as were sequences with extremely high numbers of RBD mutations relative to other sequences, leaving 679,502 retained sequences. All RBD amino-acid mutations were enumerated compared to the reference Wuhan-Hu-1 SARS-CoV-2 RBD sequence (Genbank MN908947, residues N331-T531). We acknowledge all contributors to the GISAID EpiCoV database for their sharing of sequence data (all contributors listed at: https://github.com/jbloomlab/SARS-CoV-2-RBD_MAP_Rockefeller/blob/main/data/gisaid_hcov-19_acknowledgement_table_2021_03_15.pdf).

### Data visualization

The static logo plots in the paper were created using dmslogo (https://jbloomlab.github.io/dmslogo/) version 0.6.2; a markdown rendering of the code that creates these logo plots is at https://github.com/jbloomlab/SARS-CoV-2-RBD_MAP_Rockefeller/blob/main/results/summary/escape_profiles.md. For each plasma, the y-axis is scaled to be the greatest of (a) the maximum site-wise escape metric observed for that plasma, (b) 20x the median site-wise escape fraction observed across all sites for that plasma, or (c) an absolute value of 1.0 (to appropriately scale plasma that are not “noisy” but for which no mutation has a strong effect on plasma binding).

In **Figure S4**, mutations are colored by prior deep mutational scanning measurements of yeast-displayed RBD ACE2 affinity and RBD expression from Starr et al. (2020)^32^, which are available at https://github.com/jbloomlab/SARS-CoV-2-RBD_DMS/blob/master/results/single_mut_effects/single_mut_effects.csv.

The multidimensional scaling in **Figure 1C** and **Figure 4A** that projects the antibodies into a two-dimensional space of escape mutations was performed using the Python scikit-learn package. We computed the similarity and dissimilarity in the escape maps between each pair of antibodies, then performed metric multidimensional scaling with two components on the dissimilarity matrix exactly as defined in Greaney et al. (2021)^30^. In **Figure 1C**, the multidimensional scaling shows antibodies as pie charts colored proportionally to the total squared site escape that falls into that RBD structural region. The code that generates these logo plot visualizations is available at https://github.com/jbloomlab/SARS-CoV-2-RBD_MAP_Rockefeller/blob/main/results/summary/mds_escape_profiles.md.

The interactive visualizations of the escape maps and their projections on the RBD-antibody structures available at https://jbloomlab.github.io/SARS-CoV-2-RBD_MAP_Rockefeller/ were created using dms-view (https://dms-view.github.io/docs/)^56^.

The static structural views (**Figure 2A**, **S5**) in the paper were rendered in PyMOL using antibody-bound RBD structures. PBDs are as follows: C105, 6XCN; LY-CoV016, 7C01; C110, 7K8V; C135, 7K8Z; C144, 7K90; C002, 7K8S; C121, 7K8Y. In **Figure S5B**, 7K8T is used instead of 7K8S to illustrate the quaternary antibody epitope. For identifying contact sites to highlight in Figure 1B logo plots or to classify sites in Figure 2B as contact sites (within 4A of antibody) or antibody-proximal sites within 4–8A, the following PDBs were used: 6XCM and 6XCN for C105, 7K8S and 7K8T for C002, 7K8X and 7K8Y for C121, 7K90 for C144, 7K8Z for C135, and 7K8V for C110)^25, 29^. Structural distances were computed using the bio3d package in R^57^. Surface representations of the RBD for non-antibody-bound structures utilize PDB 6M0J^58^.

In many of the visualizations, the RBD sites are categorized by epitope region (class 1, class 2, or class 3), defined by^25^ and colored accordingly. We define the class 1 epitope as residues 403+405+406+417+420+421+453+455-460+473-476+486+487+489+504, the class 2 epitope to be residues 455+456+472+483-487+489+490+491+492+493+494, and the class 3 epitope to be residues 345+346+437-452+496+498-501. There are 5 residues that overlap between the class 1 and class 2 epitopes: 455+456+486+487+489.

## Acknowledgments

We thank Andrea Loes for experimental assistance and Cathy Lin for administrative support; Dolores Covarrubias, Andy Marty, and the Genomics and Flow Cytometry core facilities at the Fred Hutchinson Cancer Research Center for experimental support; J. Vielmetter, P. Hoffman, and the Protein Expression Center in the Beckman Institute at Caltech for expression assistance. This work was supported by the NIAID / NIH (R01AI141707 and R01AI127893 to J.D.B., T32AI083203 to A.J.G., P01 AI138398-S1 to M.C.N. and P.J.B.) and the Gates Foundation (INV-004949). Support was also provided by the Caltech Merkin Institute for Translational Research (P.J.B.), a George Mason University Fast Grant (P.J.B.), and ATAC consortium EC 101003650 (D.F.R.); NIH grants U19 AI111825 and NIH U01 AI151698 for the United World Antiviral Research Network, UWARN (M.C.N., D.F.R.). The Scientific Computing Infrastructure at Fred Hutch is funded by ORIP grant S10OD028685. T.N.S. is a Washington Research Foundation Innovation Fellow at the University of Washington Institute for Protein Design and a Howard Hughes Medical Institute Fellow of the Damon Runyon Cancer Research Foundation (DRG-2381-19). C.O.B was supported by the Hanna Gray Fellowship Program from the Howard Hughes Medical Institute and the Postdoctoral Enrichment Program from the Burroughs Wellcome Fund. J.D.B., P.D.B., and M.C.N. are Investigators of the Howard Hughes Medical Institute. The content is solely the responsibility of the authors and does not necessarily represent the official views of the US government or the other sponsors.

## Author contributions

Conceptualization, A.J.G., C.O.B., M.C.N., P.J.B., and J.D.B.; Methodology, A.J.G., T.N.S., and J.D.B.; Investigation, A.J.G.; Software, A.J.G., T.N.S., and J.D.B.; Formal Analysis, A.J.G. and J.D.B.; VSV escape data, Y.W., F.S., D.P.; Resources, M.C.N., D.F.R., M.C., C.G., A.C., M.A., S.F., Z.W., F.M.; Writing – Original Draft, A.J.G. and J.D.B.; Writing – Review and Editing, all authors; Supervision, T.H., P.D.B., M.C.N., P.J.B., and J.D.B.

---

**Figure S1.**
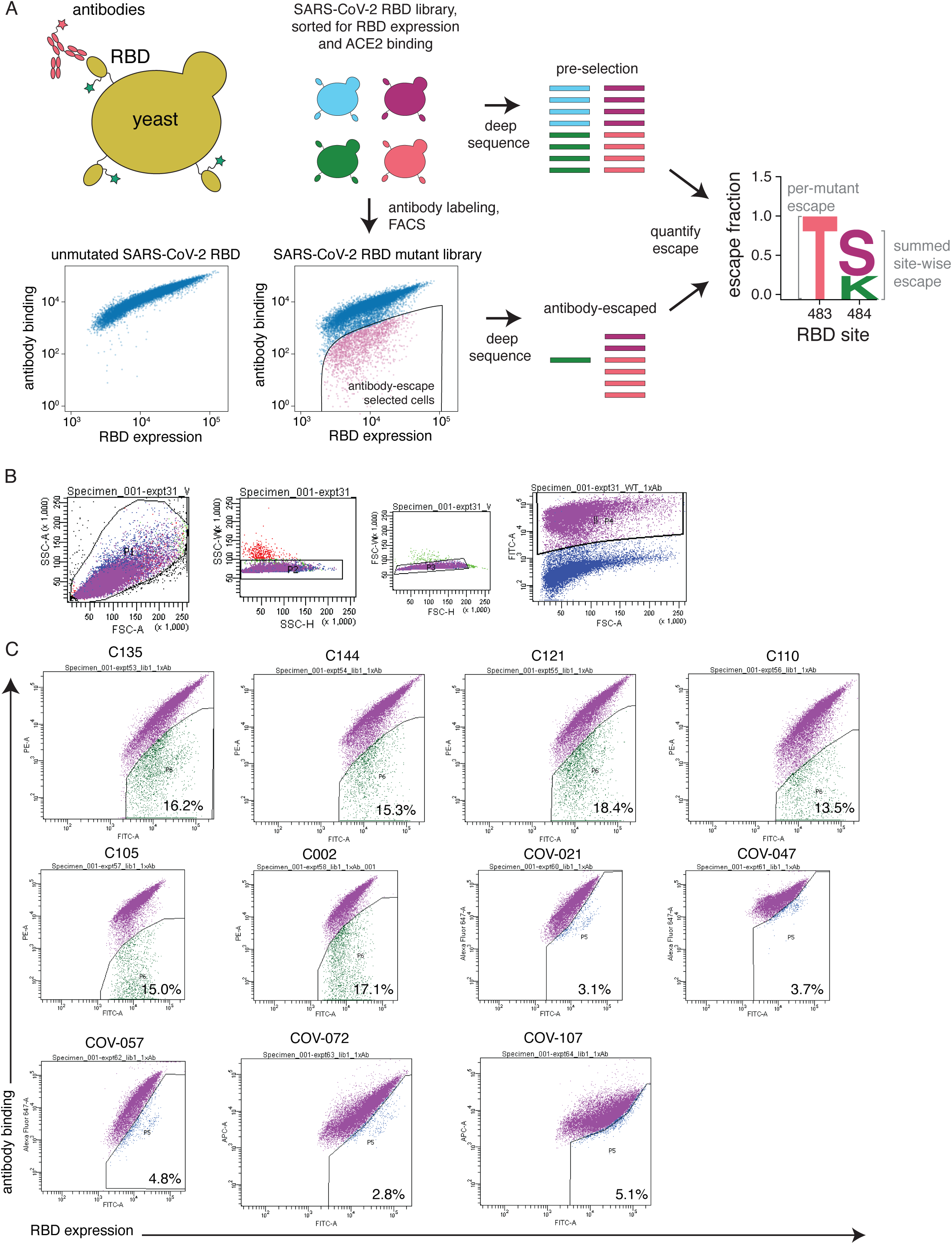
Approach for mapping RBD mutations that reduce binding by monoclonal antibodies or polyclonal plasma, related to Figure 1. **(A)** The RBD is expressed on the surface of yeast. Flow cytometry is used to quantify both RBD expression (via a C-terminal MYC tag) and antibody binding to the RBD protein expressed on the surface of each yeast cell. A library of yeast expressing different RBD mutants were incubated with antibodies or plasma and binding was detected using a IgG or IgA+IgG+IgM secondary antibody for monoclonal antibodies or polyclonal plasma, respectively. We then used FACS to enrich for cells expressing RBD that bound reduced levels of antibody, and deep sequencing to quantify the frequency of each mutation in the initial and “antibody escape” cell populations. We quantified the effect of each mutation as the “escape fraction,” which represents the fraction of cells expressing RBD with that mutation that fell in the “antibody escape” FACS bin. Escape fractions are represented in logo plots, with the height of each letter proportional to the effect of that amino-acid mutation on antibody binding. The site-level escape metric is the sum of the escape fractions of all mutations at a site. Note that both experimental and computational filtering steps were used to remove RBD mutants that were misfolded or completely unable to bind the ACE2 receptor (see **Methods**). **(B)** Representative plots of nested FACS gating strategy used for all plasma selection experiments to select for single cells (SSC-A vs. FSC-A, SSC-W vs. SSC-H, and FSC-W vs. FSC-H) that also express RBD (FITC-A vs. FSC-A). **(C)** FACS gating strategy for one of two independent libraries to select cells expressing RBD mutants with reduced binding by monoclonal antibodies or polyclonal plasma (cells in blue). Gates were set manually during sorting. Different strategies were used for monoclonal antibodies vs. polyclonal plasma. For monoclonal antibodies, selection gates were set to capture up to 95% of yeast cells expressing unmutated RBD, stained with an antibody concentration 100x lower than that used for library staining. For polyclonal plasma, selection gates were set to capture 3-6% of the RBD+ library. The same gate was set for both independent libraries stained with each plasma, and the FACS scatter plots looked qualitatively similar between the two libraries. For information on the fraction of library cells that fall into each selection gate, see **Supplementary Table 2**.

**Figure S2.**
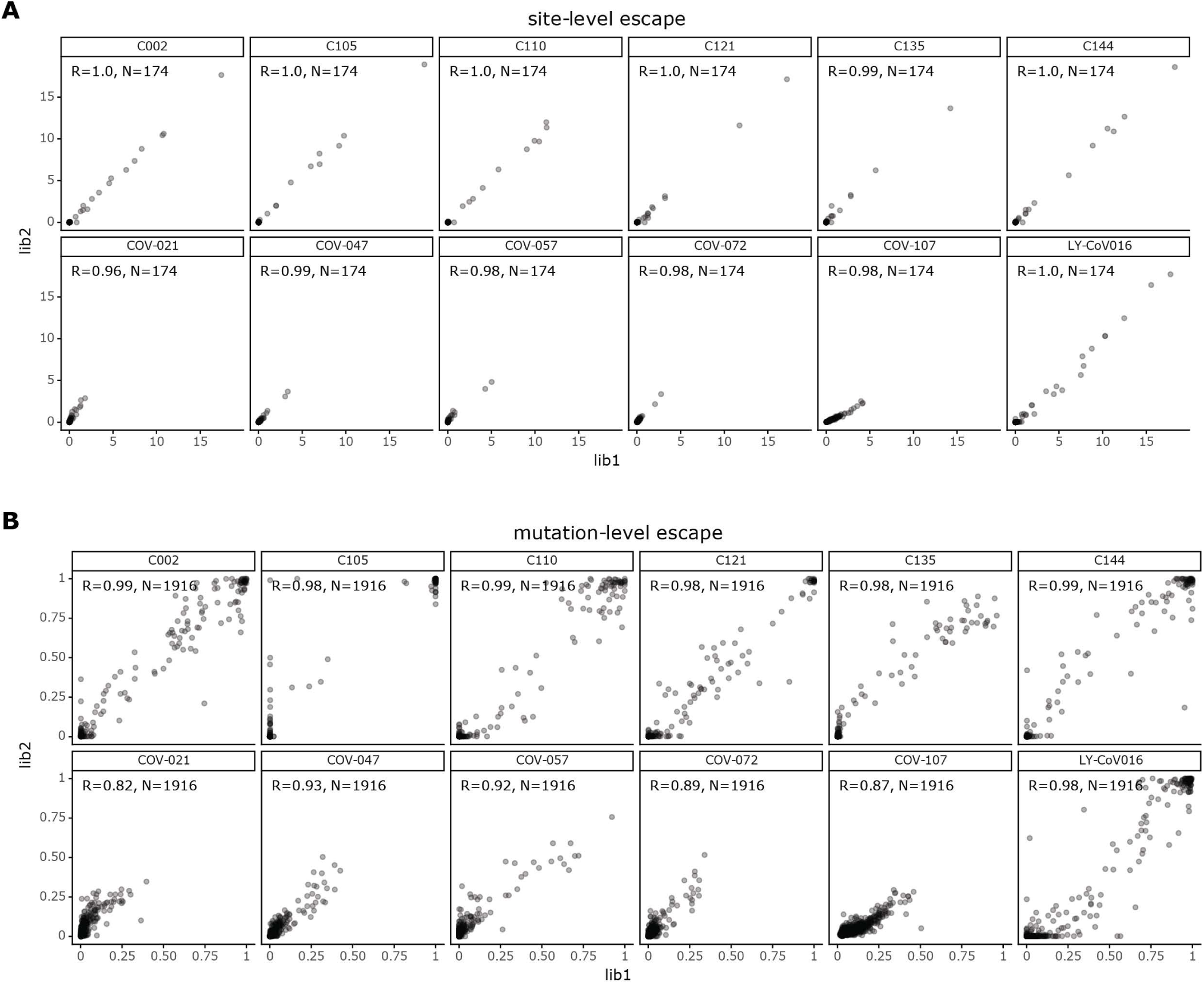
Correlations between replicates for site- and mutation-level escape metrics, related to Figure 1. **(A)** Correlation plots of site-level escape for each of the two independent RBD mutant libraries for each antibody or plasma. Each point represents one site in the RBD. **(B)** Correlation plots of mutation-level escape for each of the two independent RBD mutant libraries for each antibody or plasma. In this plot, each point represents a different mutation.

**Figure S3.**
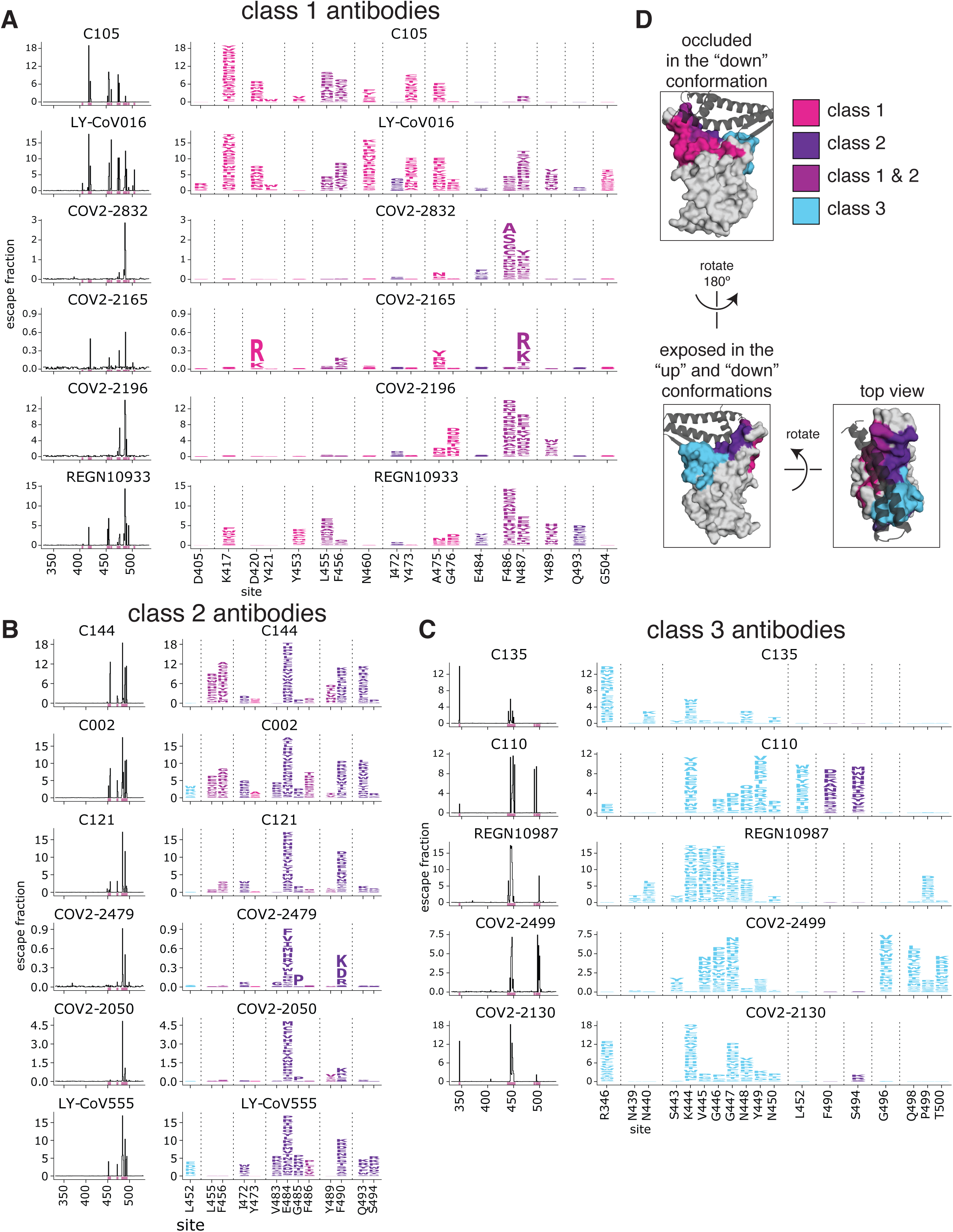
Escape maps for class 1, 2, or 3 antibodies we have profiled here or in previous studies, related to Figure 1. Escape maps for **(A)** class 1, **(B)** class 2, or **(C)** class 3 antibodies shown in Figure 1C. All escape maps were previously generated^30, 33, 35, 36^ except for C105, C144, C002, C121, C135, and C110 which are new to this study. Different sets of key sites are shown for each of the three antibody classes (see **Methods**). Sites are colored by RBD epitope as in Figure 1, also shown in panel **(D)**. For the escape maps of class 4 antibodies shown in Figure 1C, see Greaney et al. (2021)^30^.

**Figure S4.**
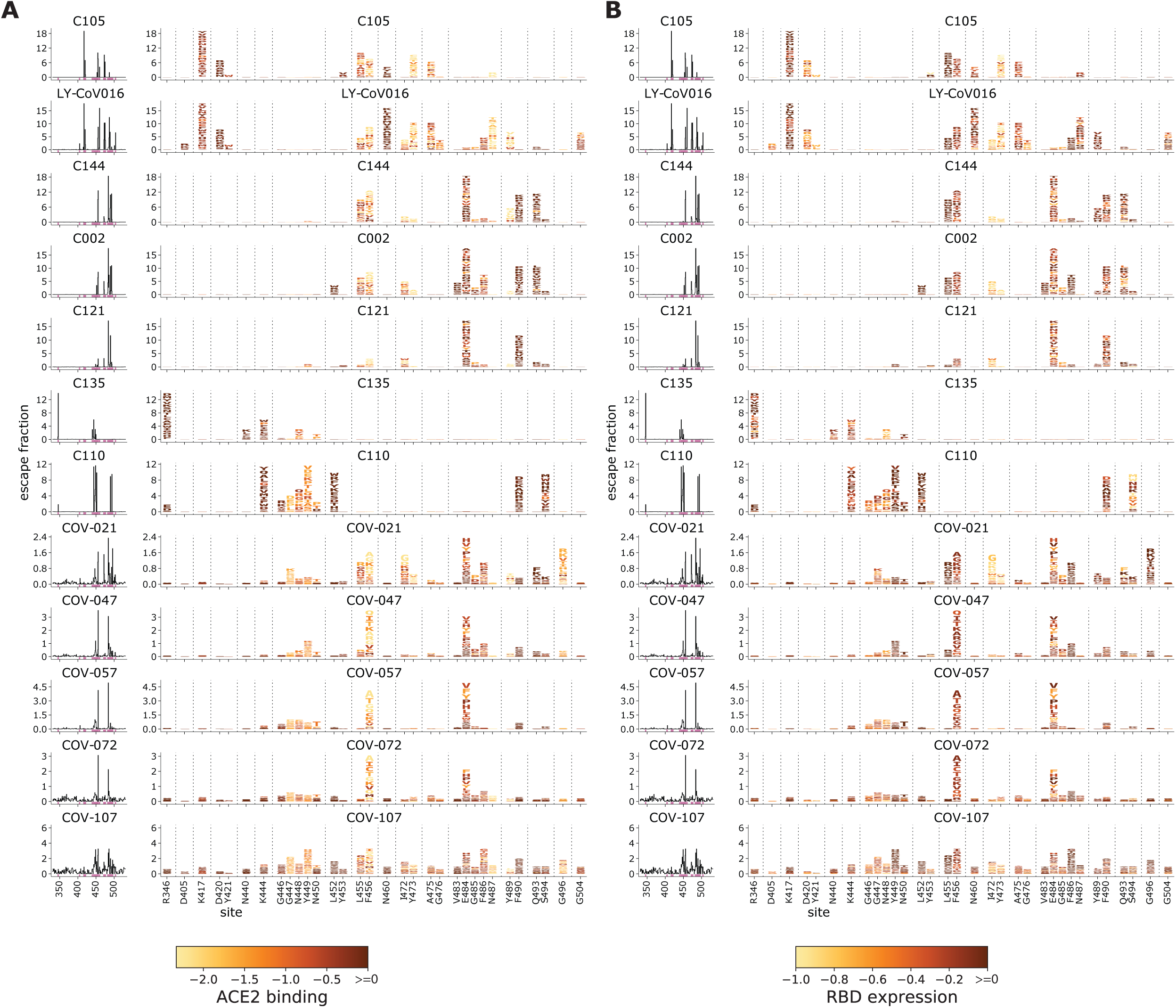
Logo plots colored according to effects of mutations on ACE2 binding and RBD expression, related to Figures 1, 3, and 4. Logo plots show the escape fractions of each mutation at key sites (any site called as a site of “strong escape” for any antibody or plasma). Letters are colored by how mutations affect RBD affinity for ACE2 or RBD expression as measured via yeast display^32^, with yellow indicating poor affinity or expression and brown indicating good affinity and expression. The top 6 rows of logo plots are for monoclonal antibodies; the bottom 5 rows are for polyclonal plasma.

**Figure S5.**
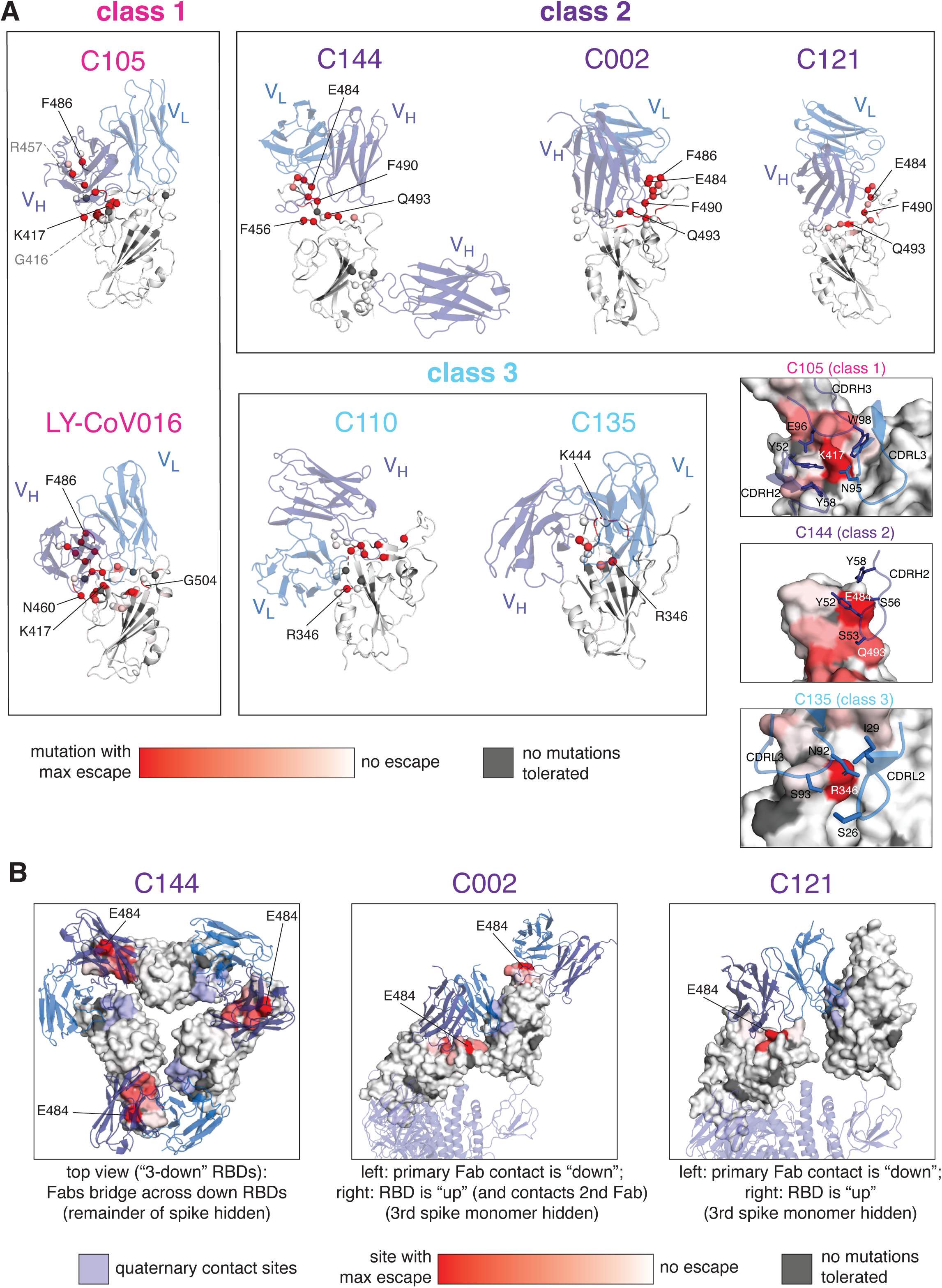
Visualization of the maximum escape at a site mapped onto cartoon representations of antibody-bound RBD, related to Figure 2. **(A)** Mapping of maximum antibody escape at a site to the antibody-bound RBD. Antibody-contact sites on the RBD (within 4Å) are shown as spheres. Sites with no escape measurements due to excessive functional constraint on the site are shown in dark gray. Each site is colored according to the maximum escape fraction of any mutation at that site (whereas Figure 2 shows site total escape), scaled from white (no escape) to red (maximum escape for any mutation for that antibody). Inset panels at right indicate key RBD-antibody interactions where mutations to the indicated RBD site disrupt antibody binding. RBD color scale indicates site total escape, as in Figure 2A. **(B)** Visualization of class 2 antibody quaternary epitopes. The total escape at each site is mapped onto the surface of the Fab-bound RBD as in Figure 2A, with white indicating no escape, and red indicating the site with the most escape from that antibody. Sites where no mutations are tolerated are indicated in dark gray. Antibody quaternary contact sites are shown in periwinkle. The C144 antibody binds to spike trimer in the “all RBDs down” conformation and forms a quaternary epitope that bridges across two adjacent RBDs by binding to a hydrophobic RBD cavity at the base of the N343 N-linked glycan. The C002 and C121 antibodies, when bound to a “down” RBD, can form a quaternary epitope with an adjacent “up” RBD. The “up” RBD also contacts another C121 Fab^25^. Our yeast-display system utilizes monomeric RBD and therefore does not map escape mutations to quaternary contact sites. These results thus cannot be used to determine the importance of the quaternary sites for antibody binding. Previous work, however, has shown that the V367F RBD mutation to the C144 quaternary epitope does not affect neutralization of pseudotyped lentivirus by C144^7^. See **Methods** for PDB accession codes used to generate structural representations.

**Figure S6.**
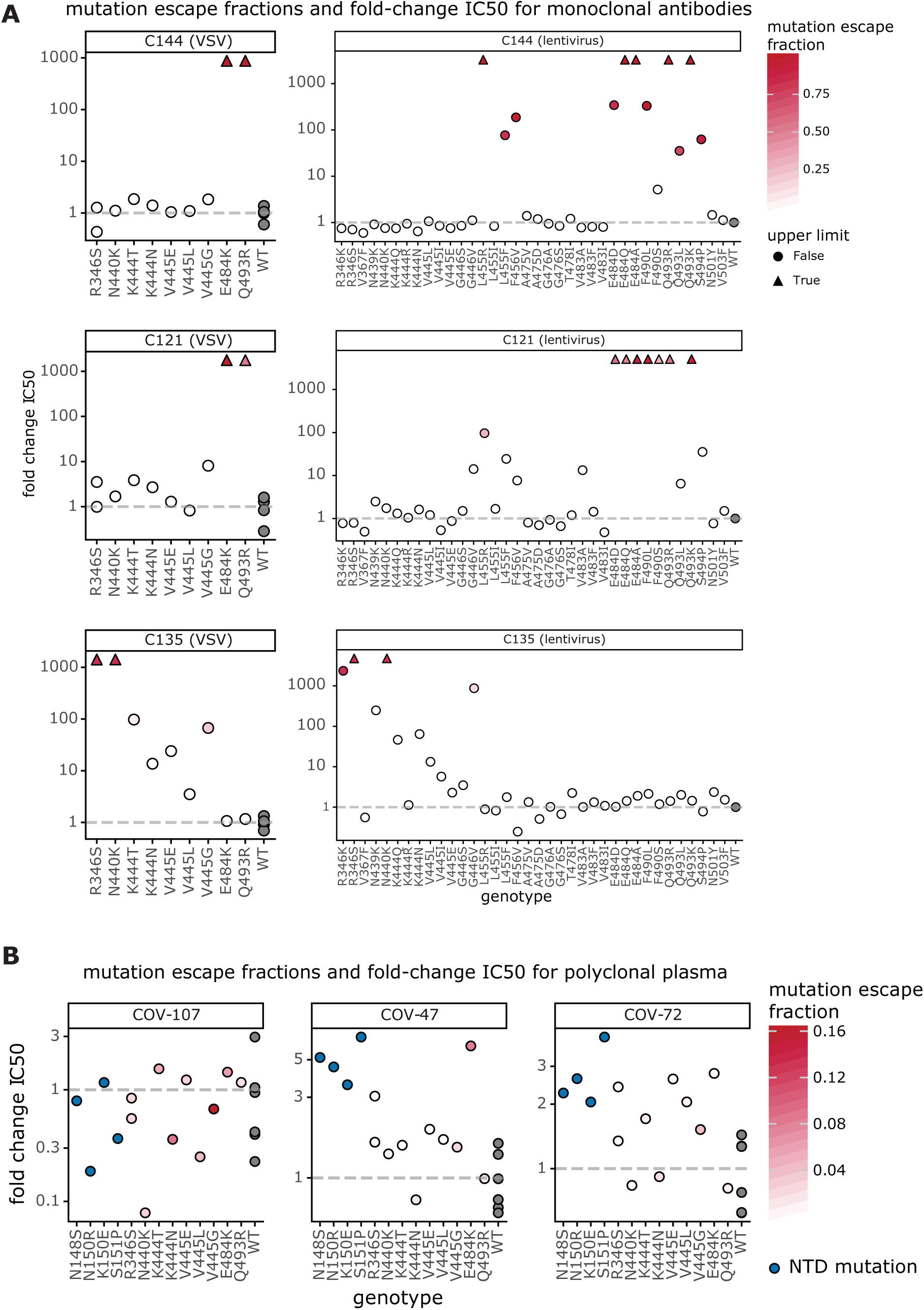
The previously measured effects of spike mutations on neutralization for 3 monoclonal antibodies and 3 polyclonal plasma, related to Figure 4. **(A)** The effects of mutations on neutralization of chimeric VSV encoding the SARS-CoV-2 (left) or spike-pseudotyped lentivirus particles (right) by monoclonal antibodies from Weisblum et al. (2020)^7^. The y-axis shows the fold-change in IC50 compared to the Wuhan-Hu-1-like spike, such that larger numbers are greater reductions in neutralization sensitivity. Mutations that had IC50s at or above the limit of detection are indicated as triangles. Points are colored according to their mutation escape fraction (Figure 1**, Supplementary Table 1**). Wildtype is in gray. **(B)** The effects of mutations on neutralization of spike-encoding chimeric VSV by polyclonal antibodies from Weisblum et al. (2020)^7^. Wildtype spike IC50 values are shown in gray. NTD mutations are shown in blue. Escape fraction color scales are independent in **(A)** and **(B)**.

## Supplemental Files

**Supplementary Table 1. Measurements of effects of all amino-acid mutations to the RBD on binding of monoclonal antibodies or polyclonal human plasma, related to Figure 1.**

The file gives the “escape fraction” for each mutation, as well as the total escape fraction at each site and the maximum escape fraction for any mutation at the site. The file is also available on GitHub at https://github.com/jbloomlab/SARS-CoV-2-RBD_MAP_Rockefeller/blob/main/results/supp_data/all_samples_raw_data.csv.

**Supplementary Table 2. Information on FACS sorting to select cells expressing RBD mutants with reduced binding by antibodies or plasma, related to Figure 1 and Figure S1.**

The file gives the number of antibody-escaped cells collected per selection for each replicate library and the percent of RBD+ cells in the antibody-escape gate for each selection, and the exact dilution used for each plasma selection. The file is also available on GitHub at https://github.com/jbloomlab/SARS-CoV-2-RBD_MAP_Rockefeller/blob/main/data/SupplementaryTable2.xlsx.

